# Invasive *Vespa velutina nigrithorax* hornets are more susceptible to entomopathogenic fungus than two other hymenopteran species, the wasp *Vespa vulgaris* and the bumblebee *Bombus terrestris*

**DOI:** 10.1101/2024.11.25.625208

**Authors:** Mathilde Lacombrade, Naïs Rocher, Blandine Mahot-Castaing, Fanny Vogelweith, Mathieu Lihoreau, Denis Thiéry

## Abstract

Entomopathogenic fungi are commonly used as biological control agents. Recently, a strain of *Metarhizium robertsii* (Sorokin, 1883) (Hypocreales: Clavicipitaceae) collected in France was identified as a good candidate for controlling invasive populations of yellow-legged hornet, *Vespa velutina nigrithorax* (Buysson, 1905) (Hymenoptera: Vespidae) in Europe. Since *M. robertsii* is generalist, its use as bioagent could be damaging for non-targeted species. Here, we compared the lethal effects of three concentrations of this *M. robertsii* strain on the invasive hornet and two non-targeted species commonly found in Europe, the common wasp *Vespa vulgaris* (Linnaeus, 1758) (Hymenoptera: Vespidae) and the buff-tailed bumblebee *Bombus terrestris* (Linnaeus, 1758) (Hymenoptera: Apidae). Exposure to *M. robertsii* spores altered the survival of all three species but at different degrees. Hornets had consistently lower survival, even under low concentrations of spores, when compared to wasps and bumblebees. Such lower susceptibility of beneficial Hymenoptera is encouraging in the perspectives of using *M. robertsii* as biocontrol agent against invasive hornets.

## Introduction

The yellow-legged hornet (*Vespa velutina nigrithorax*) (Buysson, 1905) (Hymenoptera: Vespidae) is an invasive social insect species throughout the world, especially in Europe (Haxaire et al. 2006), Korea (Kim et al. 2006) and Japan (Sakai and Takahashi 2014) (Thiery and Monceau 2024). Their geographic expansion poses major concerns to beekeeping activities and the environment (Monceau et al, 2014). Current methodologies to control hornet populations are expensive and use pesticides harmful for the biodiversity (Kishi and Goka 2017; Goulson 2019). Recently, however, several biocontrol options have been explored (Turchi and Derijard 2018) including the use of an entomopathogenic fungus, *Metarhizium robertsii* (Sorokin, 1883) (Hypocreales: Clavicipitaceae) (Poidatz et al. 2018). This fungus pertains to the complex of species *M. anisopliae* that is already commercialized for biocontrol and known to be efficient against Orthopterans, Isopterans and Thysanopterans (i.e. *M. anisopliae* strain F52 approved in Europe, USA, Brazil, Columbia and China for the use against various insects; *M. anisopliae* strain FI-985 approved in Australia, India and China for the use against grasshoppers and locusts). In particular, Poidatz et al. (2018) have demonstrated that the exposure to spores of four isolates of *M. robertsii* (EF2.5(1), EF3.5(1), EF3.5(2), EF3.5(4)) at 10^7^ sp/ml reduced by half the survival of yellow-legged hornets after only 5 to 6 days.

From a biological control perspective, entomopathogenic fungi like *M. robertsii* constitute good candidates to target a social insect such as hornets. On the one hand, sociality favors parasite transmission between genetically close individuals that are spatially aggregated, and engaged in numerous behavioral interactions, through social contacts, food exchange, or proximity with nest materials (Dimbi et al. 2013; Khun et al. 2021). However, the potential infection of non-targeted species should also carefully be taken into consideration if infected hornets leave their nest and fly, disseminating spores from their body surface (Smagghe et al. 2013). In particular, there is a risk that wild and managed pollinators get contaminated through direct contact with spores when contaminated hornets hunt unsuccessfully and release their prey (Ueno 2015), or land on a honeybee hive flight board, or contaminate flowers on which their collect nectar (Kapongo et al. 2008).

Several studies on the complex species of *M. anisopliae* (including *M. robertsii*) (Rehner and Kepler 2017) have reported negligible (Smagghe et al. 2013) to strong negative impact (Shaw et al. 2002) on several beneficial insects such as pollinators (e.g. honey bees, bumblebees), depending on the doses (Kanga et al. 2003; Smagghe et al. 2013), the formula (Ball et al. 1994), and the methods of application used (Rodríguez et al. 2009). For instance, the contact with a powder concentrate of 10^7^ sp/g decrease the survival of bumblebees by almost 25% within 6 weeks (Smagghe et al. 2013). The decrease of survival reaches 100% at a concentration of 10^9^ sp/g. The survival of honey bees was lower when the fungus was sprinkled on and between the frames of a hive (decreased by around 55%) than when conidia were stamped on filter papers and placed on half the frames (decreased by 5%) (Rodríguez et al. 2009). Given these potential advantages and disadvantages for hornet biocontrol, there is a need to better characterize the impact of *M. robertsii* exposure with comparable methodologies both on invasive hornets and non-target species before it can be further considered for biocontrol.

Here we compared the lethal effect of a *M. robertsii* strain EF3.5(1) on *V. v. nigrithorax* and two non-target pollinator species commonly found in the invaded area, the common wasp *Vespula vulgaris* (Linnaeus, 1758) (Hymenoptera: Vespidae) and the buff-tailed bumblebee *Bombus terrestris* (Linnaeus, 1758) (Hymenoptera: Apidae). We hypothesized that *M. robertsii* would have different effects depending on the ecology and the natural history of the host species. Specifically, we expected that the virulence of the parasite would be stronger in hornets than in the two other species, because these invasive insects only build aerial nests which could greatly limit the exposition to *M. robertsii*, whereas the local non-target species may have co-evolved with it since long by nesting in soil. We tested this hypothesis by exposing hornets, wasps, and bumblebees to different concentrations of *M. robertsii* and monitoring their survival for eight days.

## Methods

### Insects

All the experiments were performed in Bordeaux (France) during summers 2021 to 2023. We used four wild *V. v. nigrithorax* nests, one wild *V. vulgaris* nest, and four commercial *B. terrestris* nests (see Table 1). Although commercially reared since about 30 years, these bumblebee populations originate from wild populations sampled in Europe and are frequently mixed with freshly caught wild individuals (Velthuis and Doorn 2005). For each species, we collected the adults from the different nests and grouped them into microcolonies of 10±3 individuals in plastic boxes (72 × 26.5 × 25 cm). Each microcolony was provided sucrose solution (40% v/v) and water *ad libitum*. Bumblebee microcolonies were also supplied with a commercial mixture of pollen (Icko-apiculture company, France).

**Table 1.** Details of all the insect colonies used.

|  | Hornet | Wasp | Bumblebee |
| --- | --- | --- | --- |
| European evolution history | Invasive | Non-invasive | Non-invasive |
| Nest | Aerial | Underground<br>Occasionally aerial | Underground |
| Origin | Wild | Wild | Commercial |
| Number of nests | 4 | 1 | 4 |
| Origin<br>(Department code) | Laburgade (46)<br>Bouliac (33)<br>Bouliac (33)<br>Cahors (46) | Saucats (33) | Netherlands<br>Netherlands<br>Netherlands<br>Netherlands |
| Collection date | 2021-10-12<br>2021-10-18<br>2021-10-25<br>2021-11-28 | 2021-12-10 | 2021-06-08<br>2021-06-08<br>2021-06-08<br>2021-06-08 |
| Approximate nest size<br>(cm) | <30<br>30-80<br>30-80<br>>80 | 60-70 | <20<br><20<br><20<br><20 |
| Number of microcolonies |  |  |  |
| Control | 9 | 6 | 6 |
| 10<sup>3</sup> sp/ml | 9 |  | 6 |
| 10<sup>7</sup> sp/ml | 9 | 6 | 6 |

### Entomopathogen fungus

We used the strain EF3.5(1) of *M. robertsii* sampled in 2015 in an experimental INRAe vineyard (Villenave d’Ornon, France) (see Poidatz et al. 2018 for details). We selected this particular isolate because of its high virulence against hornets as shown by the direct inoculation and inoculation by transfer between hornets (Poidatz et al. 2018). *M. robertsii* was cultivated on a Petri dish (9 cm diameter) containing oat agar media (Oat flour 40g and Potatoes Dextrose Agar (Sigma Aldrich), Chloramphenicol (Sigma Aldrich), and permuted water) in a dark room. The fungus was transplanted every 2-3 months to ensure its healthy growth through nutrient resupplying.

### Infections

We prepared spore solutions by mixing *M. robertsii* spores (extracted from culture media), distilled water, and 0.02% Tween80 (Sigma Aldrich) (Poidatz et al. 2018; Ponchon et al. 2022). The concentration of the solution was adjusted at 10^3^ spores/ml (identified in a preliminary experiment as the sublethal concentration for bumblebees, named *low concentration*) or 10^7^ spores/ml (named *high concentration*) by counting spores in 10µl using a Neubauer cell. The control treatment was composed of distilled water and 0.02% Tween80. Each insect was exposed to one of these treatments by immersion of their entire body in the solution for 1 second. We used *M. robertsii* exposure by immersion because of its higher efficacy on hornet survival compared to exposure by contact (infected support or individual) or by food (Poidatz et al. 2018).

### Survival assays

We started the survival survey immediately after exposure to spores and the formation of microcolonies. From then on, we recorded the number of daily-dead insects and removed them from each microcolony for 8 consecutive days. We also monitored food consumption by weighing the containers of aqueous solutions (sucrose solution and water) every day.

### Statistical analysis

We ran all the analyses in R 4.3.3. We measured the effect of *M. robertsii* spore concentrations on survival for each species using a mixed-effect Cox model (*coxme* function, *survival* package, Therneau 2023) with treatment as a fixed factor and microcolony as a random effect. We ran a Cox proportional hazards regression model (*coxph* function, *survival* package, Therneau 2023) with the treatment as a fixed factor for wasps as they came from a single mother nest. We measured species sensibility differences by comparing the survival of species when exposed to a similar treatment. We used a mixed Cox model with species as a fixed factor and microcolony as a random factor. We calculated an estimate of the survival curve for each treatment within species using the *survfit* function (*survival* package, Therneau 2023) with the treatment as a fixed factor.

## Results

### Hornets were the most impacted by spore exposure

We first tested the influence of *M. robertsii* concentrations on the survival of hosts (Fig. 1). For all three species, individuals exposed to high concentrations of spores showed reduced survival compared to unexposed controls of the same species (see Table 2 for statistics). Lowly-exposed hornets showed a significantly decreased survival by 3.16 times (Cox, 95% confidence interval [CI]: 1.21-8.24, p=0.048) compared to controls. By contrast, in bumblebees, there was no difference between lowly-exposed individuals and controls (Table 2). Highly-exposed hornets showed a decreased survival by 36.83 times (Cox, CI: 14.58-93.08, p<0.001) compared to controls. This reduction was much higher than the reduction by 6.43 times (Cox, CI: 3.32-12.45) observed in wasps or by 1.61 times (Cox, CI: 0.76-3.44) in bumblebees at the same spore concentration. Overall, the difference in survival between exposure doses was larger in hornets than in wasps and in bumblebees (ANOVA, hornets: Treatment: X^2^=73.43, df=2, p<0.001; wasps: Treatment: X^2^=40.67, df=1, p<0.001; bumblebees: Treatment: X^2^=7.88, df=2, p=0.019) indicating that non-targeted species (wasps and bumblebees) were less susceptible to spore exposure than hornets.

**Table 2.**
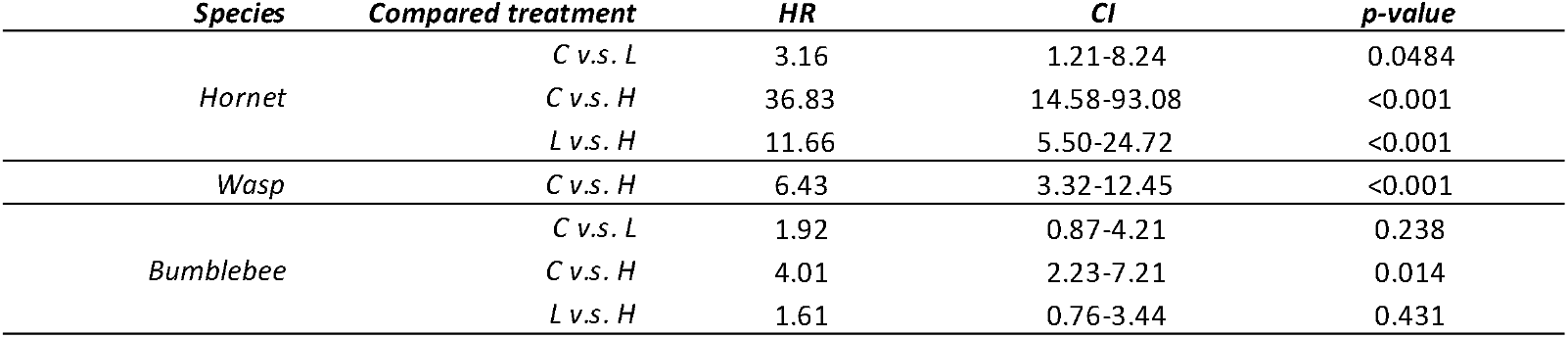
Comparison of survival probability across treatment and for each species using Cox model. Treatments: C (control), L (low spore concentration), H (high spore concentration). HR (Hazard Ratio): the risk of an event occurring (in this case, the death of an individual) when comparing one treatment with another, CI (Confidence Interval): 95% confidence interval and p-value (the probability of this risk occurring at random).

| *Species* | *Compared treatment* | *HR* | *CI* | *p-value* |
| --- | --- | --- | --- | --- |
| *Hornet* | *C v.s. L* | 3.16 | 1.21-8.24 | 0.0484 |
|  | *C v.s. H* | 36.83 | 14.58-93.08 | <0.001 |
|  | *L v.s. H* | 11.66 | 5.50-24.72 | <0.001 |
| *Wasp* | *C v.s. H* | 6.43 | 3.32-12.45 | <0.001 |
| *Bumblebee* | *C v.s. L* | 1.92 | 0.87-4.21 | 0.238 |
|  | *C v.s. H* | 4.01 | 2.23-7.21 | 0.014 |
|  | *L v.s. H* | 1.61 | 0.76-3.44 | 0.431 |

**Fig. 1:**
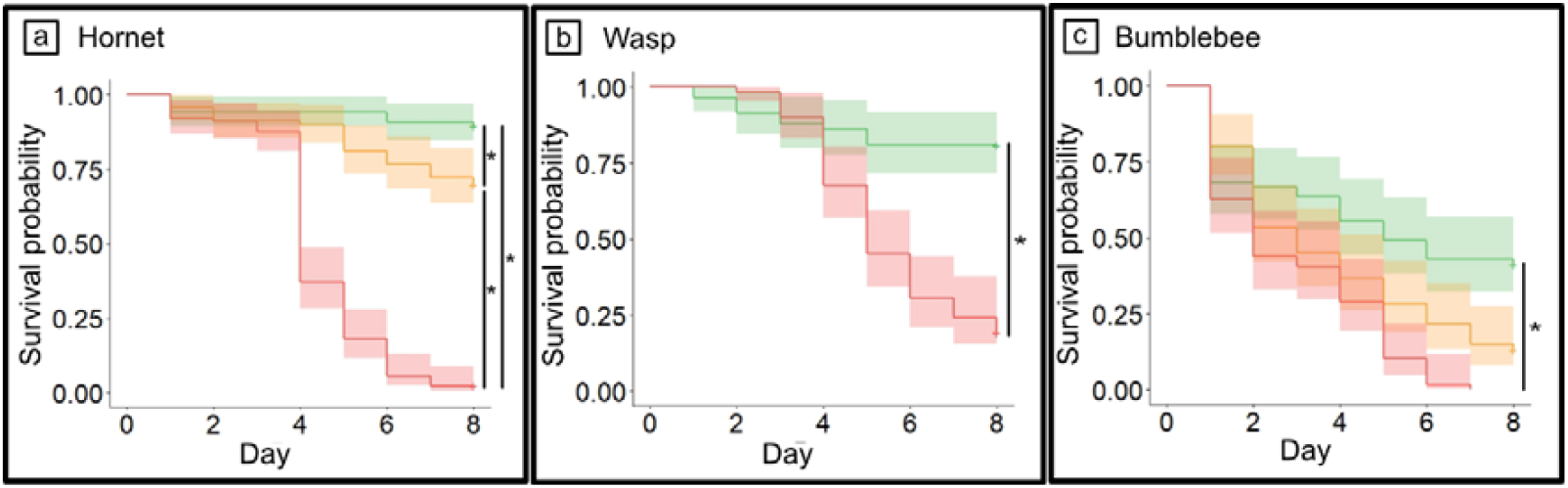
Survival probability of the three Hymenoptera tested during eight days: a) hornet, b) wasp, c) bumblebee. We exposed the insects to three treatments: control (green and upper line), low concentration of spores (10^3^ sp/ml, orange and middle line when three lines are present), high concentration of spores (10^7^ sp/ml, red and lower line). Solid lines represent experimental data and colored areas, the 95% confidence interval. Stars represent significant differences between groups calculated with a mixed-effect Cox model (a,c) or a Cox proportional hazards regression model (b).

### Hornets and wasps lived longer than bumblebees at all spore concentrations

We then compared the effects of the parasite on host survival between species and for each treatment. In unexposed controls, hornets and wasps showed a similar lifespan (Table 3 for statistics) and had a higher probability of survival than bumblebees (Table 3). When they were exposed to the parasite at low or high concentrations, hornets, and wasps also lived longer than bumblebees. At high concentrations, wasps survived longer than hornets (Table 3).

**Table 3.** Comparison of survival probability across species and for each treatment using Cox models. Treatments: C (control), L (low spore concentration), H (high spore concentration). HR (Hazard Ratio): the risk of an event occurring (in this case, the death of an individual) when comparing one species with another, CI (Confidence Interval): 95% confidence interval and p-value (the probability of this risk occurring at random).

| *Treatment* | *Compared species* | *HR* | *CI* | *p-value* |
| --- | --- | --- | --- | --- |
| *Control* | *H v.s. W* | 1.38 | 0.23-8.46 | 0.935 |
|  | *H v.s. B* | 10.62 | 2.07-54.44 | 0.0128 |
|  | *W v.s. B* | 7.69 | 1.28-46.26 | 0.0666 |
| *Low concentration* | *H v.s. B* | 8.35 | 3.08-22.67 | <0.001 |
| *High concentration* | *H v.s. W* | 0.44 | 0.25-0.72 | 0.004 |
|  | *H v.s. B* | 1.75 | 1.05-2.92 | 0.083 |
|  | *W v.s. B* | 4.13 | 2.31-7.40 | <0.001 |

## Discussion

We exposed three Hymenoptera species to different concentrations of a strain of entomopathogenic fungus recently reported to affect invasive yellow-legged hornets (Poidatz et al. 2018). Overall, for all spore concentrations tested, hornets were more susceptible to spore exposure than wasps and bumblebees, comforting the idea that *M. robertsii* is a good candidate for biocontrol against yellow-legged hornets with negligible impact on non-targeted species.

Our parasite strain EF3.5(1) harmed differently the survival of the three insect species depending on the exposure concentration. In particular, bumblebees had a shorter lifespan than the two other species but seemed less sensitive to entomopathogenic concentrations than the hornets. Since *B. terrestris* nests are found underground (Meyling and Eilenberg 2007), it is possible that they have already been in contact with closely related strains of *M. robertsii* during their evolution. Cases of natural exposure of bumblebees to *Metarhizium spp*. have already been recorded (Macfarlane et al. 1995; Schmid-Hempel 2001). In the context of host-parasite coevolution, bumblebees could have developed greater resistance to this parasite (Bérénos et al. 2009) which should less (or not) be the case for yellow-legged hornets (invasive species, aerial niche) and wasps (local species but subterranean, or aerial niche). Our observations that bumblebees are less sensitive to the parasite support this hypothesis of an evolutionary adaptation. Note however that the fact that we observed a low survival of bumblebees in our experiments could be explained by the health of the colonies we received. It would be interesting to replicate these experiments on wild bumblebees although if this bias had been present, it would have influenced our results in the direction of a greater susceptibility to the parasite.

Interestingly, the effects of *M. robertsii* were the strongest on the yellow-legged hornets. Our observation that only half of the hornets survived four days after the exposition to spores is comparable to the results in Poidatz et al. (2018), thereby confirming the efficiency of *M. robertsii* against hornets. From a biocontrol perspective, the application method of *M. robertsii* seems key to enhancing selectivity: the direct application of spores into hornets’ nests could be interesting as it allows a selective application and the reduction of potential non-targeted species contamination. Moreover, it is possible that *M. robertsii* causes a range of early post-infection sublethal effects that impair hornet activity before they die. Indeed, many parasites and pathogens can alter the behavior of insects (Gómez-Moracho et al. 2017). *M. anisopliae* can alter the reproduction of cockroaches (*Blatella germanica*) (Quesada-Moraga et al. 2004), the nutrition of fruit flies (*Drosophila melanogaster*) (Toledo-Hernández et al. 2018) and also the locomotion activity of termites (Hassan et al. 2021). It is, for instance, possible that contaminated hornets are less able to leave their nest and forage, thus greatly limiting the opportunities of contaminating non-targeted species through contact with water, flowers, or honey bee hives. However, even when intraspecific contamination occurs, if the quantity of spores deposited on the surface is lower enough, it should be harmless to non-targeted species as our results demonstrate that bumblebees and wasps were not impacted by a low concentration of spores. The sublethal effects of *M. robertsii* on hornets, their ability to spread spores in the environment, and the realistic spores’ transmission to other pollinators through contaminated surfaces should therefore be further taking into consideration to establish the safety of this biocontrol agent. Further studies are also necessary to assess the actual spores’ quantity to inject in hornets’ nests which will be dependent on the parasite transmission between nestmates. Many social insects use behavioral strategies to limit the transmission of parasites or pathogens in the colony and if so, to limit the establishment of parasites and pathogens in the nest such as grooming or reduced contact with infected individuals (Cremer et al. 2007). A preliminary experiment suggests hornets increase the duration of their grooming behavior when infected by *M. robertsii* (personal observations). These hygienic behaviors and potential others could be restrictive in an entomopathogen use context.

Our study suggests entomopathogenic fungi, and particularly *M. robertsii* strains, are good candidates for biocontrol of invasive yellow-legged hornets as the higher mortality found compared to the two non-target species reinforced its potential. However, as the effect on non-targeted species is concentration-dependent, there is a need to quantify the real risk of spores spreading in the environment by an infected individual to better assess the safety of this biological control agent on local biodiversity.

## Statements and Declarations

The authors declare no competing interests. This study was funded by the ADEME (No. 2082C0061) and the ANRT (CIFRE No. 2020_0399). MLa, FV, MLi and DT designed the study. MLa, NR, BM-C performed the experiments. MLa wrote the first draft. MLa, FV, MLi and DT revised the manuscript.

## Acknowledgments

We thank Mael Rannou for helping during experiments.

## References

Ball BV, Pye BJ, Carreck NL, et al (1994) Laboratory testing of a mycopesticide on non□target organisms: The effects of an oil formulation of Metarhizium flavoviride applied to Apis mellifera. Biocontrol Sci Techn 4:289–296. 10.1080/09583159409355337

Bérénos C, Schmid□hempel P, Mathias Wegner K (2009) Evolution of host resistance and trade□offs between virulence and transmission potential in an obligately killing parasite. J Evolution Biol 22:2049–2056. 10.1111/j.1420-9101.2009.01821.x

Cremer S, Armitage SAO, Schmid-Hempel P (2007) Social Immunity. Current Biology 17:R693– R702. 10.1016/j.cub.2007.06.008

Dimbi S, Maniania NK, Ekesi S (2013) Horizontal transmission of Metarhizium anisopliae in fruit flies and effect of fungal infection on egg laying and fertility. Insects 4:206–216. 10.3390/insects4020206

Gómez-Moracho T, Heeb P, Lihoreau M (2017) Effects of parasites and pathogens on bee cognition. Ecol Entomol 42:51–64. 10.1111/een.12434

Goulson D (2019) The insect apocalypse, and why it matters. Curr Biol 29:R967–R971. 10.1016/j.cub.2019.06.069

Hassan A, Huang Q, Mehmood N, et al (2021) Alteration of termite locomotion and allogrooming in response to infection by pathogenic fungi. J Econ Entomol 114:1256–1263. 10.1093/jee/toab071

Haxaire J, Tamisier J-P, Bouguet J-P (2006) Vespa velutina Lepeletier, 1836, une redoutable nouveauté pour la faune de France (Hym., Vespidae). Bulletin de la Société entomologique de France 111:194–194. 10.3406/bsef.2006.16309

Kanga LHB, Jones WA, James RR (2003) Field trials using the fungal pathogen, Metarhizium anisopliae (Deuteromycetes: Hyphomycetes) to control the ectoparasitic mite, Varroa destructor (Acari: Varroidae) in honey bee, Apis mellifera (Hymenoptera: Apidae) Colonies. J Econ Entomol 96:1091–1099. 10.1093/jee/96.4.1091

Kapongo JP, Shipp L, Kevan P, Sutton JC (2008) Co-vectoring of Beauveria bassiana and Clonostachys rosea by bumble bees (Bombus impatiens) for control of insect pests and suppression of grey mould in greenhouse tomato and sweet pepper. Biol Control 46:508–514. 10.1016/j.biocontrol.2008.05.008

Khun KK, Ash GJ, Stevens MM, et al (2021) Transmission of Metarhizium anisopliae and Beauveria bassiana to adults of Kuschelorhynchus macadamiae (Coleoptera: Curculionidae) from infected adults and conidiated cadavers. Sci Rep-UK 11:2188. 10.1038/s41598-021-81647-0

Kim J-K, Choi M, Moon T-Y (2006) Occurrence of Vespa velutina Lepeletier from Korea, and a revised key for Korean Vespa species (Hymenoptera: Vespidae). Entomol Res 112–115

Kishi S, Goka K (2017) Review of the invasive yellow-legged hornet, Vespa velutina nigrithorax (Hymenoptera: Vespidae), in Japan and its possible chemical control. Appl Entomol Zool 52:361–368. 10.1007/s13355-017-0506-z

Macfarlane RP, Lipa JJ, Liu HJ (1995) Bumble bee pathogens and internal enemies.

Bee World Meyling NV, Eilenberg J (2007) Ecology of the entomopathogenic fungi Beauveria bassiana and Metarhizium anisopliae in temperate agroecosystems: Potential for conservation biological control. Biol Control 43:145–155. 10.1016/j.biocontrol.2007.07.007

Poidatz J, López Plantey R, Thiéry D (2018) Indigenous strains of Beauveria and Metharizium as potential biological control agents against the invasive hornet Vespa velutina. J Invertebr Pathol 153:180–185. 10.1016/j.jip.2018.02.021

Ponchon M, Reineke A, Massot M, et al (2022) Three methods assessing the association of the endophytic entomopathogenic fungus Metarhizium robertsii with non-grafted grapevine Vitis vinifera. Microorganisms 10:2437. 10.3390/microorganisms10122437

Quesada-Moraga E, Santos-Quiros R, Valverde-Garcia P, Santiago-Alvarez C (2004) Virulence, horizontal transmission, and sublethal reproductive effects of Metarhizium anisopliae (Anamorphic fungi) on the German cockroach (Blattodea: Blattellidae). J Invertebr Pathol 87:51–58. 10.1016/j.jip.2004.07.002

Rehner SA, Kepler RM (2017) Species limits, phylogeography and reproductive mode in the Metarhizium anisopliae complex. J Invertebr Pathol 148:60–66. 10.1016/j.jip.2017.05.008

Rodríguez M, Gerding M, France A, Ceballos R (2009) Evaluation of Metarhizium anisopliae var. anisopliae Qu-M845 Isolate to control Varroa destructor (Acari: Varroidae) in laboratory and field trials. Chilean J Agric Res 69:. 10.4067/S0718-58392009000400009

Sakai Y, Takahashi J (2014) Discovery of a worker of Vespa velutina (Hymenoptera: Vespidae) from Tsushima Island, Japan. Jpn J Syst Entomol 32–36

Schmid-Hempel P (2001) On the evolutionary ecology of host–parasite interactions: addressing the question with regard to bumblebees and their parasites. Naturwissenschaften 88:147–158. 10.1007/s001140100222

Shaw KE, Davidson G, Clark SJ, et al (2002) Laboratory bioassays to assess the pathogenicity of mitosporic fungi to Varroa destructor (Acari: Mesostigmata), an ectoparasitic mite of the honeybee, Apis mellifera. Biol Control 24:266–276. 10.1016/S1049-9644(02)00029-4

Smagghe G, De Meyer L, Meeus I, Mommaerts V (2013) Safety and Acquisition Potential of Metarhizium anisopliae in Entomovectoring With Bumble Bees, Bombus terrestris. Jnl Econ Entom 106:277–282. 10.1603/EC12332

Therneau TM (2023) A package to survival analysis in R

Thiery D, Monceau K (2024) Twenty years of attempting to control the Vespa velutina invasion: will we win the battle? Entomol Gen 44:479–480. 10.1127/entomologia/2024/2731

Toledo-Hernández RA, Toledo J, Sánchez D (2018) Effect of Metarhizium anisopliae (Hypocreales: Clavicipitaceae) on food consumption and mortality in the Mexican fruit fly, Anastrepha ludens (Diptera: Tephritidae). Int J Trop Insect Sc 38:254–260. 10.1017/S1742758418000073

Turchi L, Derijard B (2018) Options for the biological and physical control of Vespa velutina nigrithorax (Hym.: Vespidae) in Europe: A review. J Appl Entomol 142:553–562. 10.1111/jen.12515

Ueno T (2015) Flower-visiting by the invasive hornet Vespa velutina nigrithorax (Hymenoptera: Vespidae). Int J Chem Environ Biol Sci 3:444–448

Velthuis HHW, Doorn A van (2005) A century of advances in bumblebee domestication and the economic and environmental aspects of its commercialization for pollination. Apidologie 37:421–451. 10.1051/apido:2006019

